# NeuroFlow: An Integrated, Cross-Platform Workflow for Mouse Brain Atlas Registration and Quantification

**DOI:** 10.64898/2026.07.15.737186

**Authors:** Aarushi Rao, Henry Oo, Can Tao, Guang-Wei Zhang

**Affiliations:** Department of Neuroscience and Anatomy, School of Medicine, Virginia Commonwealth University, Richmond, VA, USA

**Keywords:** brain atlas, Allen Mouse Brain CCFv3, image registration, signal detection, regional quantification, histology, browser-based software

## Abstract

Registration of histological sections to a reference atlas is essential for anatomical localization and region-based quantitative analysis. Although established workflows are powerful, image preparation, registration, quantification, and visualization often rely on multiple software packages, some of which require platform-specific installation or locally configured programming environments. Here, we present NeuroFlow, a browser-based workflow for quantitative analysis of mouse brain histology. NeuroFlow integrates image registration, signal detection, quantification, and visualization within a single interface and operates across major operating systems without additional software installation. It supports affine and nonlinear alignment, as well as real-time oblique reslicing of the reference atlas. All processing is performed locally in a desktop browser, without requiring a local Python environment, MATLAB installation, or associated packages and toolboxes, and without uploading images to a remote server. This design preserves user control over data and keeps intermediate results accessible for inspection and review. NeuroFlow is available at https://guangweizhang.com/tool-neuroflow.html.

## Introduction

Registration of histological images to a reference atlas is a common step in quantitative neuroscience. Placing sections from different animals in a shared coordinate framework allows measurements to be assigned to the same anatomical regions and compared across animals and experiments. For mouse-brain studies, the Allen Mouse Brain Common Coordinate Framework version 3 (CCFv3) is a widely used reference atlas. It provides a three-dimensional anatomical template, standardized coordinates, and a hierarchical annotation of brain regions for spatial localization and regional analysis [1]. Earlier Allen Institute resources included genome-wide gene expression mapping [2] and the mesoscale connectome and its neuroinformatics [3–4]. Related atlas efforts include enhanced anatomical labeling and transcriptomic and cell-type atlases [5–7]. The CCFv3 anatomical template was assembled from serial two-photon tomography imaging [1, 8]. The CCFv3 now underpins large-scale studies of cortical and thalamic connectivity, single-neuron reconstruction, and cell-type characterization at regional and whole-brain scales [9–11]. Many interactive and automated approaches have therefore been developed to align mouse brain images with CCFv3.

Existing tools address different components of this problem. QuickNII and the QUINT/Nutil workflow support registration and quantification of serial rodent brain sections [12, 14–15] using volumetric reference atlases such as Waxholm Space [13]. Image-analysis platforms and libraries, including QuPath, CellProfiler, Fiji/ImageJ, ilastik, scikit-image, ITK-SNAP, and 3D Slicer, collectively support annotation, measurement, segmentation, and scripting [16–23]. Dedicated methods provide trainable pixel classification, star-convex detection, generalist cellular segmentation, U-Net–based segmentation, and large-scale image reconstruction [24–28]; a broader review provides methodological context [29]. Atlas-aware tools and workflows support one or more of atlas access, whole-brain registration, anatomical annotation, visualization, and automated cell or feature detection [30–40]. General-purpose registration toolboxes such as elastix and ANTs, together with multimodal microscopy-to-MRI pipelines, underpin many of these alignment approaches [41–43]. Meanwhile, web-based resources such as the Scalable Brain Atlas and ImJoy demonstrate that atlas access and image analysis can be delivered through a browser [44–45]. In practice, however, a complete analysis often requires users to move among several software environments. Each transfer of images, metadata, and intermediate results creates an additional opportunity for section identities, registration settings, or analysis parameters to become disconnected from the final output. This fragmentation also increases the effort required to train new users and establish reproducible workflows.

NeuroFlow addresses this fragmentation by integrating the complete workflow within a single browser-based environment. Built around the Allen Mouse Brain Common Coordinate Framework version 3 (CCFv3) [1], it combines manual initial plane assignment with affine registration, nonlinear warping, and interactive three-dimensional oblique atlas reslicing. The same application supports user-defined tissue masking, signal detection, assignment of detected signals to anatomical regions, region-specific quantitative analysis, and three-dimensional visualization. All computation is performed locally in the browser, without requiring a local Python or MATLAB environment or the upload of images to a remote server. Project states and intermediate results can be saved for subsequent inspection and review. The initial release prioritizes a complete, user-guided workflow spanning image preparation, atlas registration, quantification, and three-dimensional visualization, with an emphasis on cross-platform accessibility, ease of use, and minimal hardware and software requirements. Future releases will focus on increasing automation while preserving user control and the ability to inspect intermediate results.

## Results

### Overview of the NeuroFlow workflow from brain sections to three-dimensional quantification

NeuroFlow organizes the analysis of mouse brain histological sections cut in the coronal plane or at oblique angles to it into six stages (**Figure 1**). Cropper converts whole-slide images into editable individual section crops and can be skipped when sections have already been prepared (**Figure 1A**). Arranger establishes the order of sections while preserving section identity and allows the series to be inspected before registration (**Figure 1B**). Scaler aligns each section to the Allen Mouse Brain Common Coordinate Framework version 3 (CCFv3) [1] using affine and nonlinear transformations and supports interactive oblique atlas reslicing, with the transformation for each section saved for subsequent analysis (**Figure 1C**). Quantifier generates tissue masks and detects user-defined signals (**Figure 1D**). Analyzer assigns detected signals to anatomical regions and aggregates them into section- and series-level regional measurements and summaries (**Figure 1E**), whereas Visualizer displays the registered sections and detected signals within an interactive three-dimensional brain model (**Figure 1F**). Together, these stages provide an integrated workflow for section preparation, atlas registration, signal detection, anatomical assignment, regional quantification, and three-dimensional visualization.

**Figure 1.**
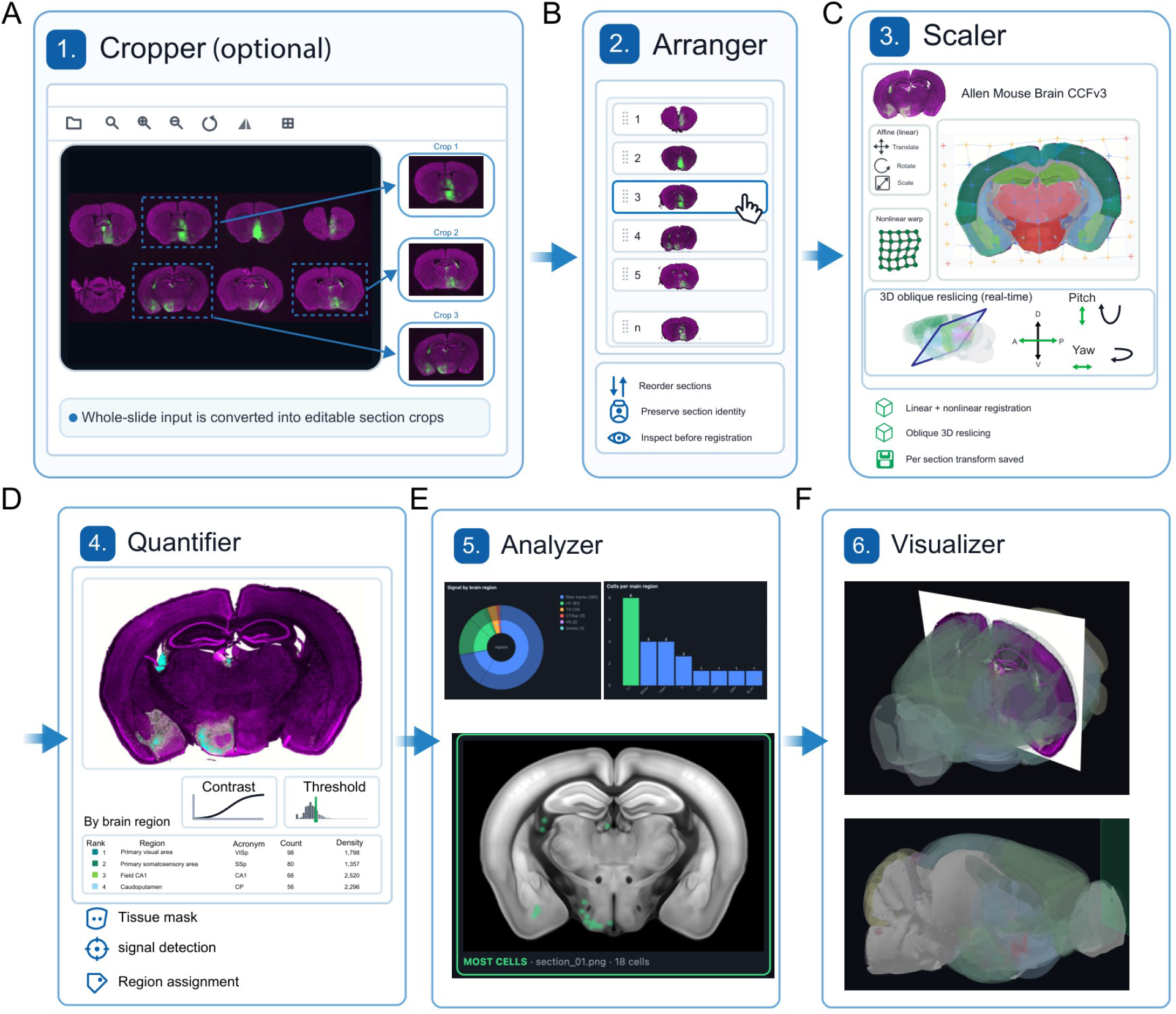
Overview of the NeuroFlow pipeline. Six linked stages are shown: Cropper, Arranger, Scaler, Quantifier, Analyzer, and Visualizer. They cover section preparation, anatomical ordering and coarse plane assignment, registration, selected-color signal detection, regional and series-level aggregation, and 3D inspection.

NeuroFlow runs locally in a desktop browser and stores intermediate project state on the user’s computer, allowing analyses to be inspected, revised, saved, exported, and reopened without uploading images to a remote server. At each stage, user-defined parameters are presented alongside the corresponding outputs. Intermediate outputs include cropped section images, section order and identity, atlas-plane assignments, affine and nonlinear transformations, tissue masks, signal maps, atlas-region assignments, regional measurements, series-level summaries, and three-dimensional visualization scenes. The project file preserves section identities, ordering, registration parameters, masks, and section-level results. However, some settings in Quantifier, Analyzer, and Visualizer are not yet serialized and are recorded separately when a complete analysis record is required.

### Optional preparation of multisection images and coarse atlas assignment

To demonstrate these optional preparation steps, we used a whole-slide histological image containing eight brain sections from a wild-type mouse that received a retrograde AAV-GFP injection (see Methods). When several sections are present in one image, Cropper allows the user to draw a separate, editable region of interest around each sample and save each region as an individual section image (**Figure 2A**). The number of regions is not fixed, and Cropper can be skipped when sections have already been saved as separate images.

**Figure 2.**
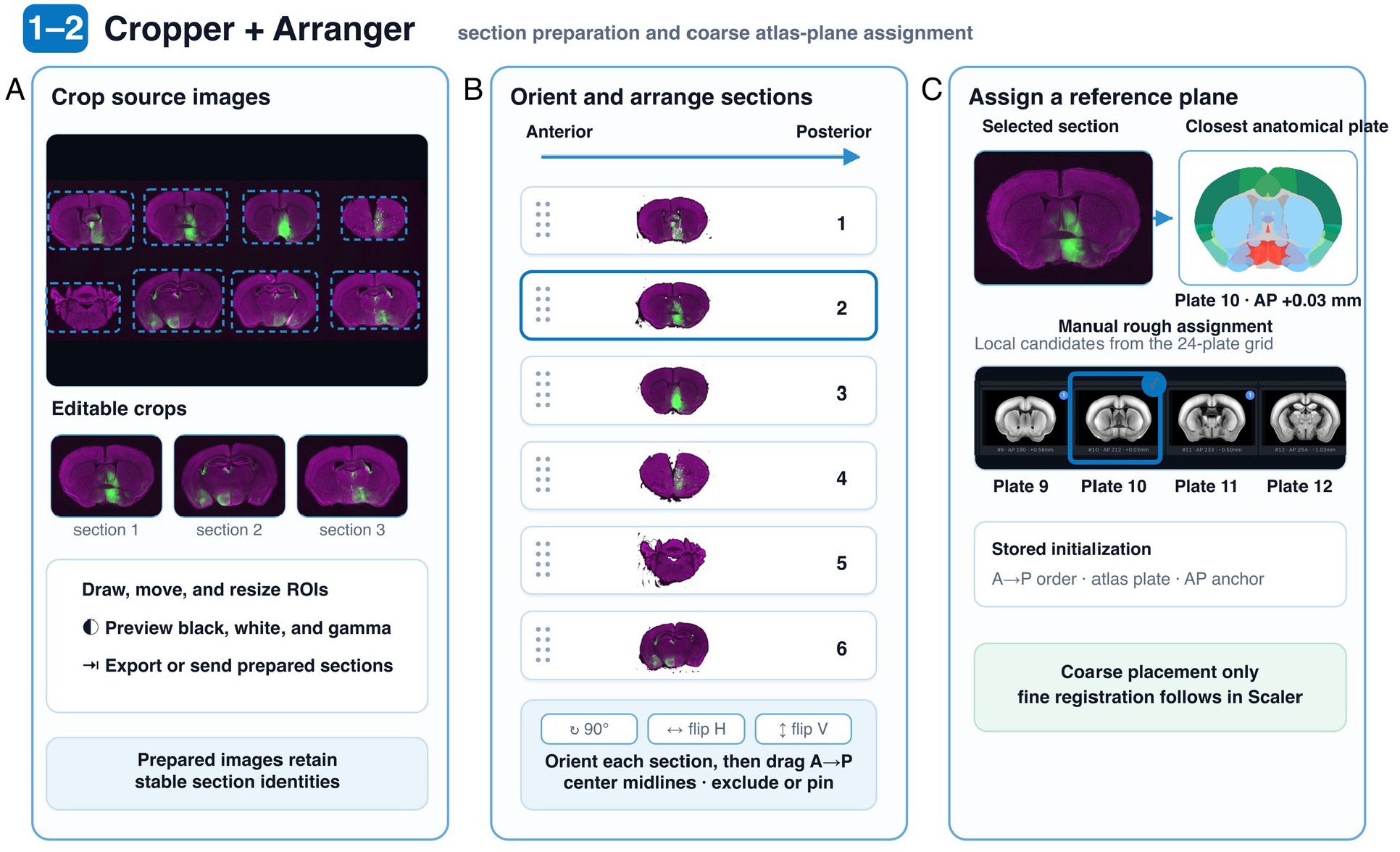
Image preparation, arrangement, and coarse atlas-plane assignment. **(A)** The Cropper divides a multi-section source image into editable regions while retaining section identity. **(B)** The Arranger provides orientation controls and direct anterior–posterior ordering. **(C)** The closest anatomical match is chosen manually from a 24-plate Allen Mouse Brain CCFv3 grid. Plate 10 (AP +0.03 mm) is the rough match shown here, and the enlarged reference is the rainbow rendering of that same resource.

Arranger can be used to correct section orientation and, if desired, place the images in anterior-to-posterior order while preserving their identities (**Figure 2B**). This step is also optional because approximate atlas assignments can establish the anatomical sequence later. The 24 representative CCFv3 plates serve as visual landmarks for manual similarity assessment; NeuroFlow does not automatically compare the section against all 24 plates. For example, the section shown in **Figure 2C** is assigned to Plate 10 (AP +0.03 mm). This coarse assignment provides a starting point for detailed affine and nonlinear registration in Scaler; it is not the final anatomical match.

### Scaler refines atlas alignment through oblique, affine, and nonlinear adjustments

Once the individual section images are ready, the next step is to align each image to CCFv3 using the tissue outline and visible anatomical landmarks. Scaler supports this step by separating the three-dimensional cutting-plane orientation from deformation within the two-dimensional tissue image. The coarse plate assignment supplies an initial reference level (**Figure 3A**). For an obliquely cut section, the user adjusts AP position, pitch, and yaw while Scaler updates the corresponding atlas reslice in real time (**Figure 3B**). Displaying the cutting plane and resliced atlas side by side makes the plane selection directly visible.

**Figure 3.**
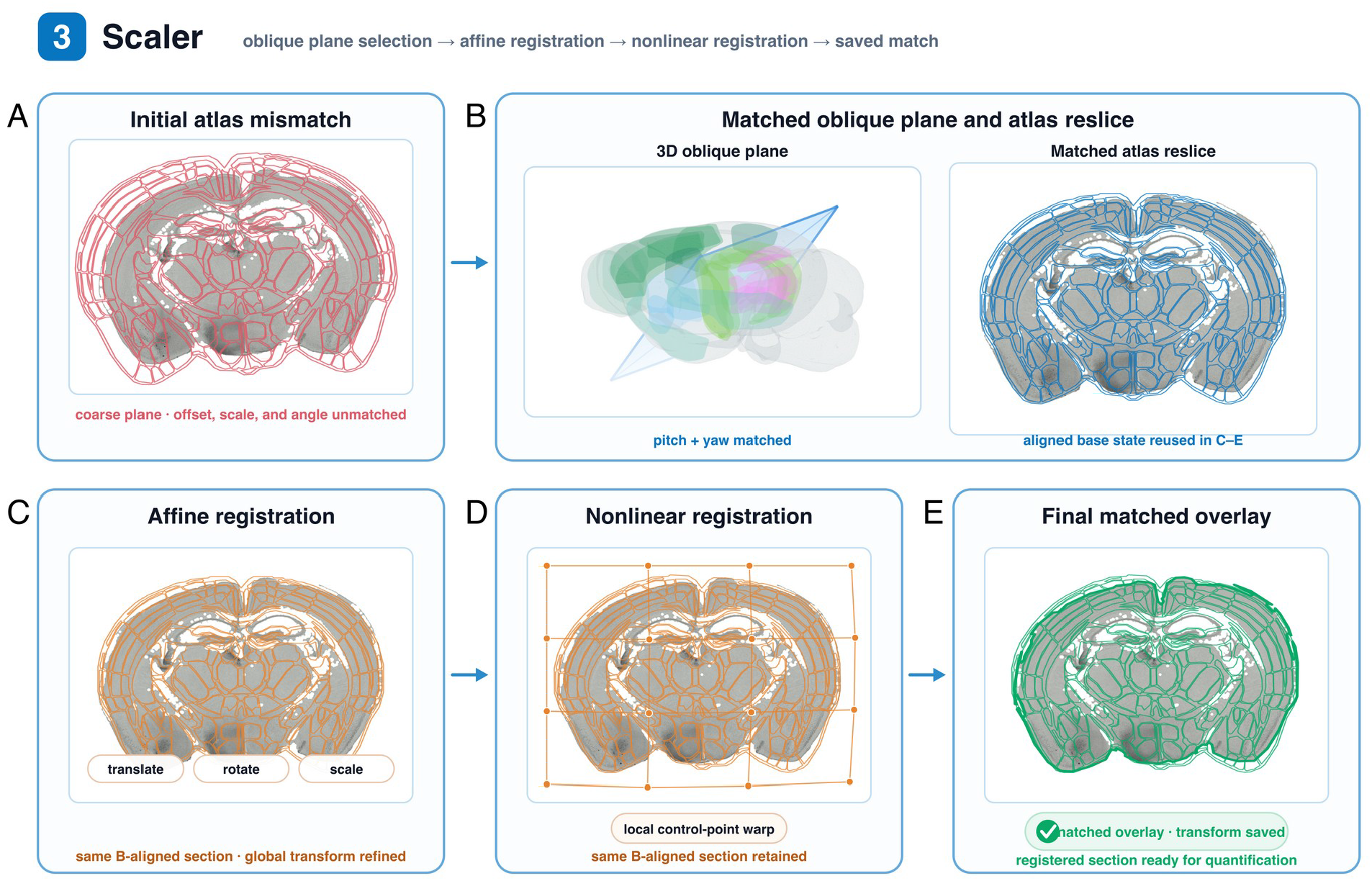
Linear and nonlinear registration with 3D oblique atlas reslicing. **(A)** The section is shown against a mismatched coarse atlas plate. **(B)** AP position, pitch, and yaw are adjusted in the enlarged 3D view, with the resulting atlas reslice displayed beside it. **(C)** Using the section aligned in panel B, affine refinement corrects translation, rotation, and x- and y-axis scale. **(D)** Starting from the affine result in panel C, nonlinear control points refine local alignment while preserving the same section and oblique plane. **(E)** The final overlay follows the tissue boundary, after which the section transform is saved.

After the atlas plane has been selected, translation, rotation, and independent horizontal and vertical scaling establish the affine, or linear, match (**Figure 3C**). Remaining local differences are corrected by dragging control points on the nonlinear grid until the atlas boundaries follow the tissue contour and internal landmarks (**Figure 3D**). The user can move freely between the linear and nonlinear views while refining the registration. The completed overlay is then reviewed, and its transform is retained for downstream regional assignment (**Figure 3E**).

### Quantifier detects and segments signal within the registered section

After registration, the next step is to detect and segment signals within the aligned tissue image. For this demonstration, Quantifier detects GFP fluorescence within an analysis region drawn directly on the section with a freehand brush; lasso and erase controls allow the region to be refined (**Figure 4A,B**). The signal threshold is freely adjustable, and the signal overlay updates with the selected setting so that detected pixels can be checked against the original microscopy image (**Figure 4C**).

**Figure 4.**
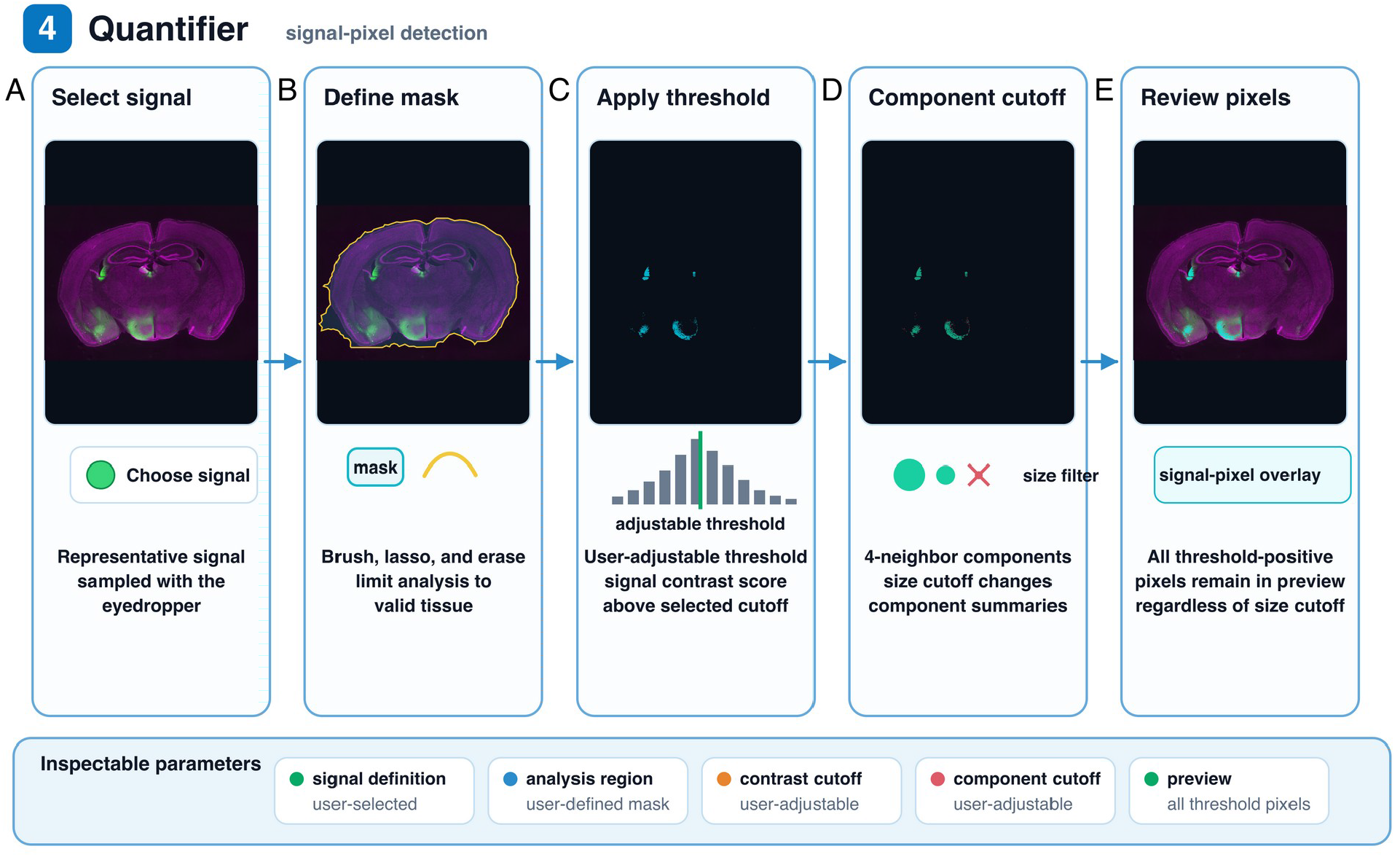
Signal-pixel detection. **(A)** A representative signal color is entered or sampled with the eyedropper; green fluorescent signals are shown here. **(B)** Brush, lasso, and erase tools define the analysis area. **(C)** A threshold is applied for signal detection. **(D)** Four-neighbor connectivity groups threshold-positive pixels, and an optional minimum size filters the component summary. **(E)** The signal preview continues to show every threshold-positive pixel regardless of that cutoff.

When component summaries are useful, adjacent positive pixels can be grouped using four-neighbor connectivity. The minimum component size is set by the user and can be adjusted to suit the signal; it acts only as a filter for the component summary (**Figure 4D**). It does not remove positive pixels from the source-pixel map, which remains visible in the signal preview (**Figure 4E**). The output therefore reports measured image pixels rather than inferred cells or synthetic centroids.

### Analyzer maps registered signals across the Allen anatomical hierarchy

After signal detection, the next step is to determine where the segmented signals lie within the registered atlas. Analyzer displays the signal-positive source pixels over the aligned CCFv3 boundaries, allowing their anatomical distribution to be inspected without the underlying microscopy intensity (**Figure 5A,B**). It then samples the registered annotation at each signal location to assign an anatomical identity (**Figure 5C**). For example, the displayed section shows how signal locations are translated into atlas-based regional measurements; the same mapping can be applied to any registered section.

**Figure 5.**
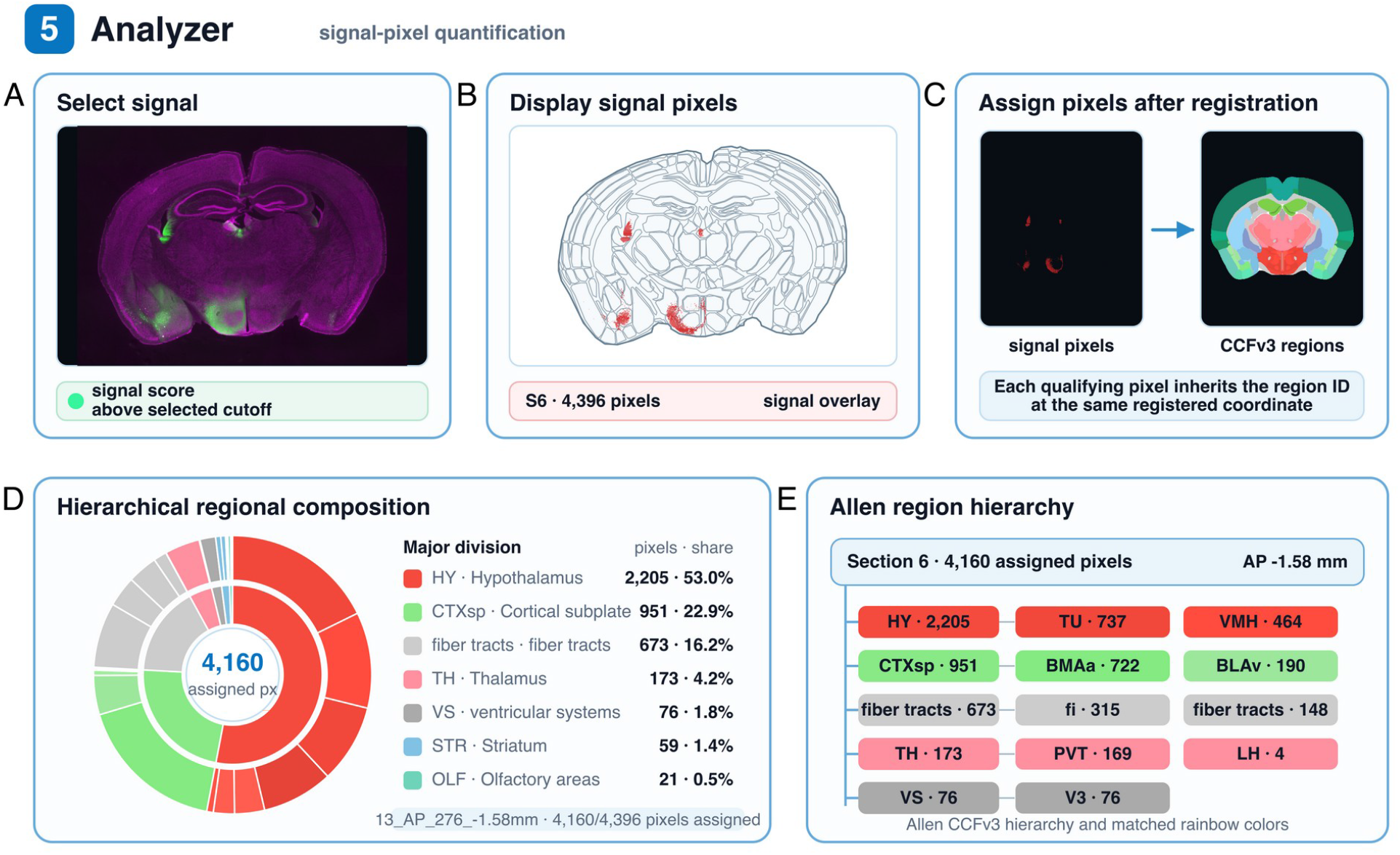
Analyzer signal-pixel quantification and Allen anatomical hierarchy. **(A)** The example image shows green fluorescent signals. **(B)** Qualifying pixels are shown as a high-contrast overlay on line-only registered CCFv3 boundaries; the microscopy intensity image is hidden. The example section contains 4,396 qualifying source pixels. **(C)** Regional identity is assigned by sampling the registered CCFv3 label image at each qualifying crop coordinate. The matched resource is Plate 13 (AP −1.58 mm), and 4,160 of 4,396 pixels fall within named Allen regions. **(D)** A two-level sunburst divides the measured count among major divisions and constituent regions. **(E)** A root-to-major-to-region tree shows the largest regional contributions. Colors in panels D and E come from the matching Plate 13 rainbow atlas.

Once assigned, the segmented signal can be quantified across the Allen anatomical hierarchy. Analyzer summarizes the same measurements from broad divisions through nested subregions, allowing the user to move from an overview to progressively finer anatomical detail (**Figure 5D**). A linked hierarchy view shows how each measurement is distributed among parent and child regions (**Figure 5E**), with colors matched to the corresponding CCFv3 rainbow atlas. The output therefore provides multilevel regional quantification rather than a flat list. These values describe segmented image signal assigned to each region and should not be interpreted as inferred cell counts.

### Visualizer places registered sections and signals within the three-dimensional brain

Finally, after regional assignment and quantification, the signals and their anatomical locations can be inspected in three dimensions. Visualizer places each registered histological plane within the exterior CCFv3 brain surface using its saved AP position, pitch, yaw, and affine parameters (**Figure 6A**). A tissue boundary derived from the purple structural channel removes the slide background and clips the section to the intersection between its plane and the three-dimensional brain surface. Internal atlas-region meshes are hidden so that the registered section remains visible. Brain Signals mode displays the segmented source pixels on the same clipped plane and uses the same camera orientation as the section view (**Figure 6B**). The linked two-dimensional view retains the original section and signal context (**Figure 6C**), while the three-dimensional scene can be rotated, panned, zoomed, and inspected interactively. This provides a spatial quality-control view of section placement and signal location. The current three-dimensional placement uses the saved oblique and affine parameters but does not apply the nonlinear control-grid warp; regional quantification therefore remains based on the registered two-dimensional annotation.

**Figure 6.**
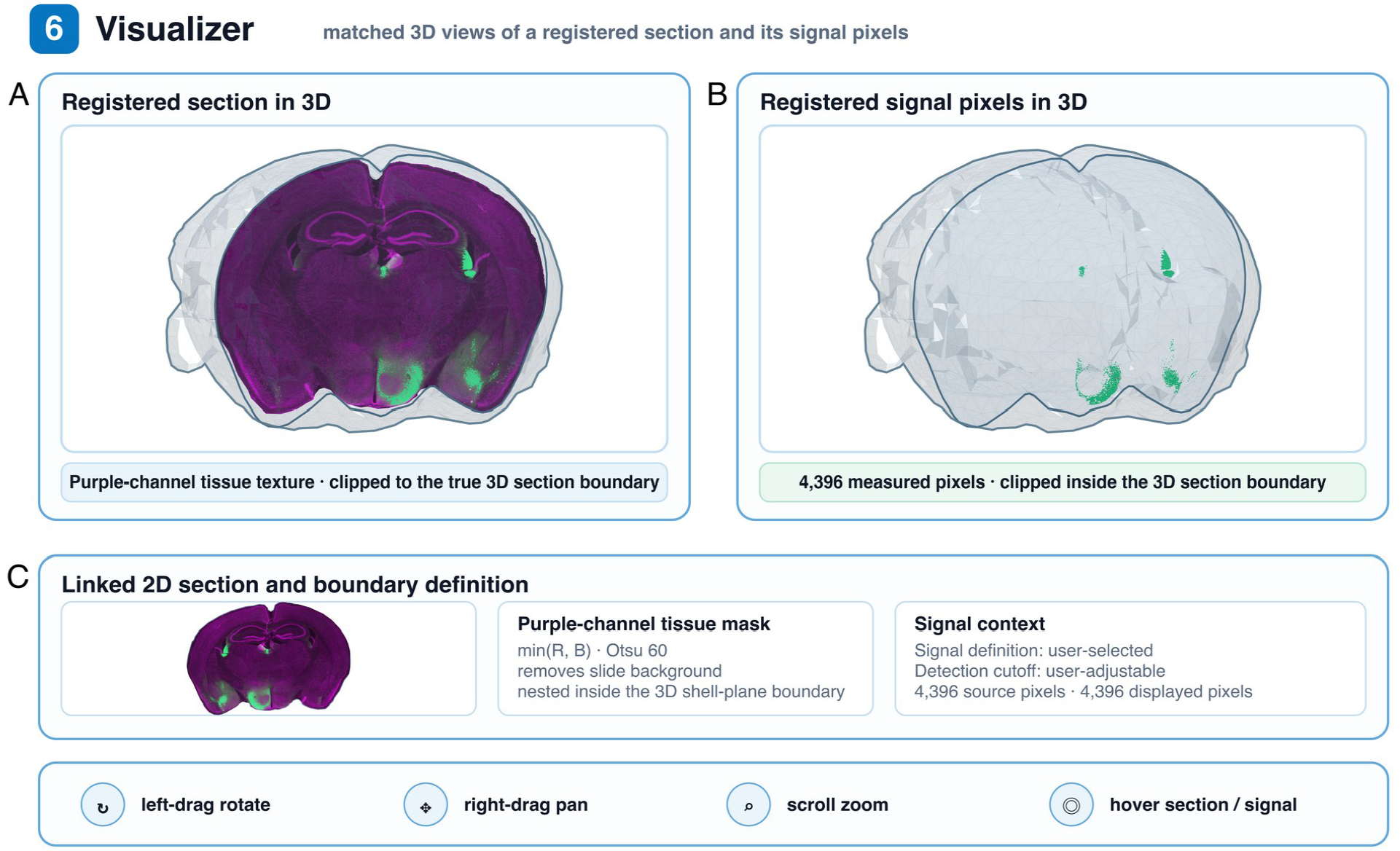
Visualizer 3D inspection of a registered section and measured signal pixels. **(A)** Slide background is removed with a purple-channel tissue mask defined as the largest filled component after Otsu thresholding of min(R, B). The section boundary is the true 3D intersection between the registered microscopy plane and the Allen CCFv3 whole-brain surface. Internal atlas meshes are hidden, and panels A and B share one oblique camera orientation. **(B)** The same view as in panel A shows only the registered segmented signals within the 3D brain contour. **(C)** A linked 2D section retains the full signal context. Users can rotate, pan, zoom, and inspect by hovering.

## Discussion

NeuroFlow integrates image preparation, section ordering, coarse atlas-plane assignment, two- and three-dimensional registration, signal detection, regional quantification, and visualization within a single browser-based workflow. The interface is designed for ease of use: source images, controls, and intermediate results remain available for inspection at each stage, while section identity and registration information stay linked to downstream measurements. Established serial-section pipelines such as QuickNII and the QUINT/Nutil workflow [12, 14–15] address registration and quantification effectively but are typically deployed as separate desktop applications. Keeping these steps together in one browser-based environment reduces the need to move files and metadata between separate applications.

Within the same application, users can crop and orient sections, choose an initial atlas plane, refine oblique, affine, and nonlinear alignment, define analysis regions, detect signals, assign them to CCFv3 annotations, examine hierarchical regional summaries, and inspect registered sections and signals in three dimensions. Processing runs locally in a desktop browser without uploading images or requiring a local Python or MATLAB environment. The first release emphasizes a lightweight, cross-platform workflow with minimal hardware demands and external software dependencies.

In the current version, all anatomical judgments are made manually and remain under the user’s control. The user selects the nearest reference plane, adjusts the three-dimensional cutting plane, refines in-plane registration, defines the analysis mask, and reviews the detection settings. Because these decisions are manual, users may introduce errors in section order, plane selection, registration, or segmentation. The relevant images and overlays remain visible so that each decision can be checked before regional aggregation. Future versions will add automation for plane selection, registration, and other repeated steps, drawing on existing automated and deep-learning approaches to section registration and cell detection [35–37, 39]. Those functions will require greater computational resources, whereas the first release is intentionally lightweight and broadly accessible across operating systems.

This release is configured for mouse brain analysis using the Allen Mouse Brain CCFv3. The same framework can be adapted to other species by supplying compatible template and annotation volumes and coordinate metadata, as provided for example by the Waxholm Space atlases of the rat brain [13, 46]; an appropriate three-dimensional surface mesh would also need to be generated or sourced separately. Quantifier and Analyzer provide complementary signal measurements: Quantifier applies a user-defined analysis mask and detection settings, whereas Analyzer evaluates registered signal pixels across the section for atlas-based regional summaries. The three-dimensional Visualizer applies the saved oblique and affine parameters; regional quantification remains based on the registered two-dimensional annotation because the nonlinear control-grid warp is not applied in the current three-dimensional view. Signal measurements represent segmented image pixels and are not automatically equivalent to validated cell counts.

## Methods

### Software architecture

NeuroFlow is a browser-based, single-page JavaScript application reached through the public NeuroFlow page. React provides the interface. Canvas 2D operations and typed arrays handle atlas rendering, registration, and oblique reslicing; the 3D Visualizer uses Three.js/WebGL. TIFF files are decoded in the browser, and tabular results can be exported as CSV or XLSX. All analysis runs on the client without a backend server. Project state is stored locally and can be saved to or restored from a project file. It includes sections, order, atlas anchors, registration parameters, mask blobs, and retained results, but not every global Quantifier setting, Analyzer display option, or Visualizer state. The six tabs are Cropper, Arranger, Scaler, Quantifier, Analyzer, and Visualizer.

### Input data and cropping

The Cropper accepts PNG, JPEG, TIFF, and OME-TIFF files. Each source image is decoded to an HTML canvas, where tissue sections are marked with axis-aligned rectangular regions of interest. The regions can be moved or resized while the image is panned or zoomed. Black-level, white-level, and gamma controls alter the display; nondefault values are incorporated into the rasterized crops passed downstream. TIFF samples are converted to 8-bit display data. The OME-TIFF reader uses the first image file directory (IFD), except when a file contains exactly three equal-sized, single-sample IFDs that can be interpreted as RGB channels. Every crop retains a persistent section identity. Orientation is corrected later in the Arranger.

### Section ordering and atlas assignment

The Arranger presents extracted sections as cards. A card records arbitrary rotation, including ±90° shortcuts, horizontal and vertical flips, exclusion, and pinning. Users drag the cards into anterior–posterior order and align the tissue midlines. Each section is then manually matched to the closest of 24 representative Allen Mouse Brain CCFv3 plates to initialize registration. The reference index and AP anchor are stored with the ordered series and can be exported as JSON or QuickNII-style XML. This assignment is only a coarse starting point; it does not perform the final registration.

### Registration and oblique reslicing

NeuroFlow includes the Allen Mouse Brain CCFv3 anatomical template and annotation data [1]. Plate metadata and the logical CCF coordinates follow a 25-µm grid of 456 ML × 320 DV × 528 AP voxels. The volumes used for interactive 3D reslicing are downsampled to 50 µm (228 × 160 × 264 voxels), and a bundled region-name table resolves annotation identifiers. In the Scaler, the user selects a section, refines its AP level, and views the atlas as a line, gray, rainbow, or combined overlay at adjustable opacity. Global registration includes translation, rotation, and separate horizontal and vertical scaling. A free-form grid with 4–30 divisions supplies nonlinear refinement; each cell is triangulated and mapped piecewise affinely. The AP level, linear parameters, and nonlinear grid are stored for each section. The matching annotation plane is regenerated when needed rather than saved in the project file.

For sections that depart from the canonical coronal plane, the Scaler samples an oblique plane through the 3D atlas volume. The plane is defined by its central AP coordinate, pitch, and yaw, with pitch and yaw adjustable over ±25°. For normalized output coordinates (x, y), the AP coordinate on the 25-µm CCF grid is a₀ + tan(yaw) × (x − 1/2) × 456 + tan(pitch) × (y − 1/2) × 320, where a₀ is the AP coordinate at the plane center. ML, DV, and AP coordinates are divided by the volume step of 2 to index the 50-µm volume. Template and annotation volumes are sampled by nearest neighbor on the same plane, keeping the displayed reference and label map synchronized. Saved in-plane affine and nonlinear transformations are then applied within the resliced plane.

### Signal detection

The Quantifier analyzes only pixels inside a user-defined mask edited with brush, lasso, erase, and pan tools. Users define the signal of interest manually or sample a representative signal with an eyedropper. A user-controlled cutoff is applied to the resulting signal-contrast score for each pixel. Pixels above the threshold appear in the signal overlay. Four-neighbor components can be summarized with an optional minimum-size cutoff, but that setting does not change the signal-pixel preview. The output is not treated as a validated cell detector.

### Regional signal-pixel quantification

Analyzer Signal mode recalculates signal-positive pixels from each full crop using the selected color preset and a configured contrast cutoff. It does not inherit the Quantifier mask, threshold, or minimum-component size. Qualifying pixels retain their crop coordinates, and the registered CCFv3 label image is sampled at those same positions. Signal-positive area, measured as qualifying-pixel count, can then be summarized by region and arranged within the Allen hierarchy. Section-level and hierarchical counts remain linked to the registered atlas identity and Allen atlas display colors. The color preset and resulting summaries are recorded separately because the current multi-sheet XLSX export omits the Analyzer Signal display.

### 3D visualization and inspection

The Visualizer displays saved registration results in a Three.js/WebGL scene, in the spirit of 3D neuroanatomical and biological-image viewers [47–48]. Each histological plane is positioned from its saved AP coordinate, pitch, yaw, translation, rotation, and x/y scale. The scene uses the Allen Mouse Brain CCFv3 whole-brain surface mesh, with internal region meshes hidden. A purple-channel tissue mask—min(R, B), Otsu threshold [49], largest filled component—first clips the plane and its signal pixels. The fitted intersection with the exterior volume provides a second clip. Brain Signals mode places the complete qualifying Analyzer pixel mask on the registered in-brain plane. Section and signal views use one oblique camera orientation. Users can rotate, pan, or zoom the scene and inspect the linked 2D context. Current 3D placement does not apply the nonlinear control-grid warp.

### Animal procedures and microscopy

All animal procedures were reviewed and approved by the Virginia Commonwealth University Institutional Animal Care and Use Committee (IACUC) and were conducted in accordance with the Guide for the Care and Use of Laboratory Animals, the Public Health Service Policy on Humane Care and Use of Laboratory Animals, and applicable institutional, state, and federal regulations. Controlled substances were handled under the responsible investigator’s current DEA registration and in accordance with VCU policy and applicable state and federal requirements. A 2.5-month-old female C57BL/6J mouse underwent stereotaxic surgery. General anesthesia was induced with 5% inhaled isoflurane and maintained at 1–2% using a calibrated R540 Enhanced Small Animal Anesthesia Machine (RWD Life Science). Body temperature was maintained using a ThermoStar Homeothermic Monitoring System (RWD Life Science). Ethiqa XR (buprenorphine extended-release injectable suspension, 1.3 mg/mL; Fidelis Animal Health) was administered at the start of surgery for perioperative analgesia according to the IACUC-approved regimen.

An 80-nL volume of pENN.AAV.EF1a.eGFP.WPRE.rBG (AAV retrograde [50]; Addgene viral prep #105547-AAVrg; RRID:Addgene_105547), a gift from James M. Wilson, was delivered stereotaxically to a basal forebrain target spanning the ventral portion of the medial septum and the anterior hypothalamic region. The recorded coordinates were AP +466 µm, ML −533 µm, and an injection depth of 4,736 µm. Virus was delivered with a nanoliter injector through a glass micropipette made from World Precision Instruments capillary tubing (WPI, Sarasota, FL, USA) and pulled on a P-97 Flaming/Brown Micropipette Puller (Sutter Instrument, Novato, CA, USA). Tissue was sectioned at 50 µm and labeled with NeuroTrace fluorescent Nissl stain (Thermo Fisher Scientific). Entire sections were imaged on a BZ-X1000 Series All-in-One Fluorescence Microscope (KEYENCE, Itasca, IL, USA) at 2× magnification using multicolor acquisition and image stitching.

Figures 4–6 were generated from these experimental histological images. Figure 4 demonstrates signal detection in a GFP-labeled section using a Quantifier contrast threshold of 50 and an eight-pixel minimum component size for summaries; the signal preview still contains every threshold-positive pixel. Figure 5 applies the Analyzer signal-detection settings with a cutoff of 40. The displayed section contains 4,396 qualifying source pixels measured directly from the microscopy image, with no inferred cells or synthetic centroids. Of these, 4,160 map to named regions on Plate 13 (AP −1.58 mm) and are grouped by the Allen hierarchy using colors from the corresponding rainbow plate. The other 236 remain in the source-pixel total but have no named Plate 13 label. Figure 6 shows all 4,396 measured pixels within the registered purple-channel tissue boundary and clipped whole-brain section. These data document the workflow and are not presented as a powered biological comparison.

### Author contributions

Aarushi Rao wrote the code and developed the tool. Henry Oo performed the viral injection, tissue sectioning, and microscopy imaging and assisted with tool testing. Can Tao and Guang-Wei Zhang designed and supervised the project. Can Tao and Guang-Wei Zhang wrote the manuscript. All authors read and approved the final manuscript.

## Acknowledgments

A.R. is supported by the Wright Center Summer Undergraduate Research Program (WSURP) at VCU. This project is supported by the Alzheimer’s Association (AARF-23-1148428), the Virginia Commonwealth University Convergence Team Planning Grant, and Virginia Commonwealth University startup funding awarded to G.-W.Z.

## Competing interests

The authors declare no competing interests.

## Data and code availability

The NeuroFlow release and study page are available at https://guangweizhang.com/tool-neuroflow.html.

## Notes

https://guangweizhang.com/tool-neuroflow.html

### Competing Interest Statement

The authors have declared no competing interest.

